# *Valeriana officinalis* genome sequence reveals candidate genes for valerenic acid biosynthesis and flavonoid metabolism

**DOI:** 10.64898/2026.08.14.744958

**Authors:** Julie Anne Vieira Salgado de Oliveira, Mariana Baez, Boas Pucker

## Abstract

*Valeriana officinalis* is the scientific name for valerian, a plant known for producing valerenic acid, a compound with anxiolytic properties. Anxiety disorders represent a significant global health crisis, impacting everyday lives. As the global demand for natural, non-synthetic anxiety treatments rises, *V. officinalis* has emerged as a promising, yet underutilized, medicinal resource. Understanding its genome is the first step toward unraveling the biosynthetic genes underlying valerenic acid production, facilitating further research into its production. Here, we report the first genome sequence of valerian, with an assembly size of 3.3 Gbp and an N50 of 110.8 Mbp, and its corresponding annotation with 96.6% completeness, providing a foundational resource for studying the genetic basis of specialized metabolism in valerian. The value of this genome sequence for discoveries in specialized metabolism is demonstrated by the identification of the flavonoid biosynthesis gene repertoire and the selection of strong candidate genes for valerenic acid biosynthesis. This genome sequence holds the potential to support future functional studies aimed at elucidating the regulation of medically relevant metabolite pathways in *V. officinalis*.

## Introduction

*Valeriana officinalis* (**Figure 1A**), also known as valerian, is a herbaceous plant in the order Dipsacales, family Caprifoliaceae, subfamily Valerianaceae, native to both Europe and Asia. Its height ranges between 1 meter and 1.5 meters, with compound or divided leaves pinnately arranged; the flowers are mostly white to pink and occur on terminal branches (**Figure 1B**). The species has an interesting evolutionary history, with diploid (2x=14), tetraploid, and octoploid plants in natural populations [1]. According to the C value database, the genome size of *V. officinalis* can vary from 1.49 pg to 4.08 pg for the 1C value (approximately 2.5 Gbp to 3.9 Gbp) [2], which reflects this underlying ploidy variation.

**Figure 1.**
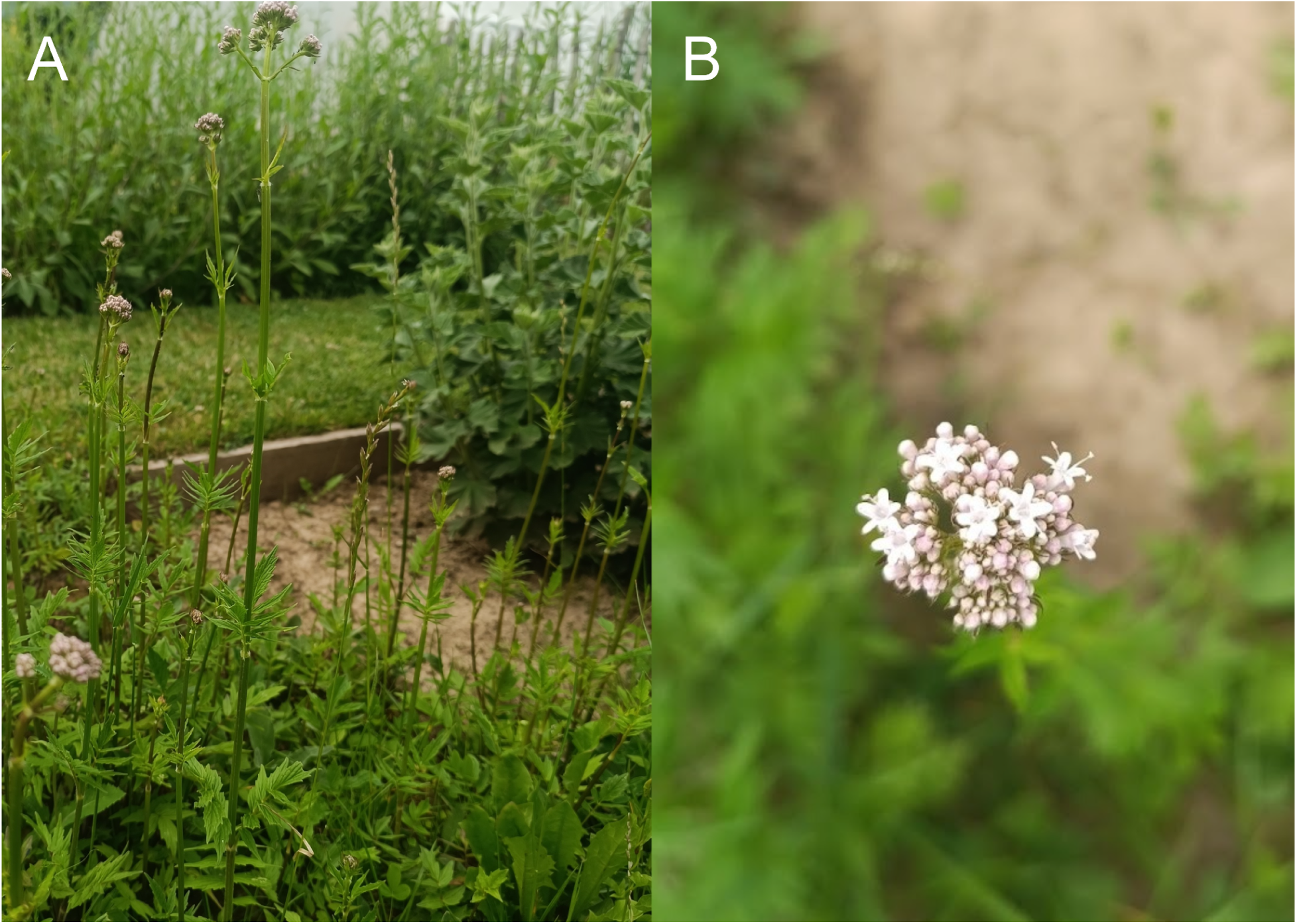
**A**: *Valeriana officinalis*, Bonn University Botanic Gardens. **B**: Close-up of *V. officinalis* flowers.

Valerian has a long history of traditional use as a mild sedative and anxiolytic, and root preparations are widely used as herbal remedies for sleep and anxiety-related problems [3,4]. Anxiety disorders represent a substantial global health burden, affecting a large proportion of the population [5], and valerenic acid, an extract from valerian roots, can be used as an adjunct or alternative to conventional pharmacotherapy. Diverse clinical and pharmacological studies have investigated the efficacy and mechanism of valerian extracts in modulating sleep quality and anxiety-related symptoms, with activity attributed primarily to sesquiterpenoids, most notably valerenic acid and its derivatives [6–8].

Valerenic acid biosynthesis begins with the cyclization of farnesyl diphosphate (FPP) by valerena-4,7(11)-diene synthase (also reported as valerena-1,10-diene synthase), the first committed and rate-limiting step of the pathway [9]. The resulting sesquiterpene is oxidized to valerenic acid through the combined activity of the cytochrome P450 VoCYP71DJ1, an alcohol dehydrogenase, and an aldehyde dehydrogenase, as demonstrated by heterologous reconstruction in yeast [10]. Terpene synthases (TPSs) are generally characterized by a conserved two-domain architecture, an N-terminal domain and a metal-binding C-terminal domain, and typically form multigene families of several dozen to over a hundred members per angiosperm genome, classified into clades TPS-a through TPS-h [11,12].

The biosynthesis of valerenic acid has been substantially elucidated through heterologous expression studies; however, the native genomic organization, copy number, and regulation of these biosynthetic genes in *V. officinalis* remain unresolved. Beyond this sesquiterpenoid pathway, valerian also accumulates flavonoids [13], indicating that the species’ specialized metabolism spans both the terpenoid and phenylpropanoid-derived branches. Chemical analyses of *Valeriana officinalis* roots have consistently reported the presence of flavonols, including quercetin and kaempferol derivatives, confirming that the flavonol branch of the pathway is active in this species [14].

Flavonoids are a diverse group of specialized metabolites that are derived from phenylalanine. The flavonoid biosynthesis pathway branches into the flavone, flavonols, proanthocyanidin, and anthocyanin biosynthesis [15]. Anthocyanins are well-known floral pigments that confer a diverse range of colors, including orange, red, pink, purple, and blue [16]. Due to their antioxidant properties, anthocyanins are considered beneficial nutritional components for humans, making them a central target of plant biotechnology applications [17,18]. A variety of modification enzymes add sugar moieties, acyl groups, or methyl groups to anthocyanins, resulting in a plethora of chemically different derivatives [16]. While anthocyanin biosynthesis has been considered well conserved across most land plants, recent discoveries of deep loss, rewiring, or substitution with another pigment biosynthesis pathway call this belief into question [19]. These recent insights into the complex evolution of pigment biosynthesis underscore the importance of investigating lineage-specific differences in specialized metabolic biosynthetic networks. A genome sequence of valerian is expected to facilitate the identification of the native gene complement underlying these pathways, including paralogs and regulatory elements that cannot be inferred from heterologous reconstruction alone.

Here, we present a genome sequence of *V. officinalis*, along with its structural and functional annotations. The flavonoid biosynthesis gene repertoire was identified, and candidate genes for valerenic acid biosynthesis were selected, including homologs of the characterized enzyme VoTPS1, thereby providing a foundational genomic resource for future functional characterization of both pathways and demonstrating the value of this genome sequence.

## Material and Methods

### Plant material and chromosome counting

*Valeriana officinalis* seeds were obtained from the Botanical Garden of TU Braunschweig (XX-0-BRAUN-7422401). The seeds were sown in August 2025 in a soil substrate, and plants were grown under long-day conditions (16 h light, 8 h darkness) in a plant cultivation room at approximately 20 °C. Since valerian can exhibit varying ploidy levels, and knowing this beforehand helps determine the amount of data to be obtained through sequencing, a chromosome count was performed.

Mitotic chromosome preparations were made from actively growing root tips collected from five individual plants in soil, treated with 8-hydroxyquinoline (2 mM) at room temperature (∼25 °C) for two hours and at 6 °C for three hours, and fixed in ethanol:acetic acid (3:1 v/v). Fixed root tips were washed in distilled water and digested in 20 µl of an enzymatic mixture (1 % pectolyase Y-23 and 2 % cellulase “Onozuka” R-10, in citric buffer) for 60 min at 37 °C, washed in 70 % ethanol, and the root section was carefully macerated with a needle. From this root solution, 8 µl was dropped onto a glass slide, fixed in ethanol: acetic acid, air-dried at room temperature, and stained with DAPI Vectashield®. At least 10 mitotic metaphases per individual were captured and analyzed using an Axiocam 305 color camera coupled to an Axiolab 5 fluorescence microscope and ZEN Blue imaging software. The images were adjusted and optimized for brightness and contrast using Adobe Photoshop 27.5.0 (2026).

### DNA extraction and Nanopore sequencing

The genome sequencing followed a previously described workflow [20]. In brief, the selected plant was incubated in the dark for one day to reduce the starch content before harvesting samples. Young leaves were harvested for high-molecular-weight DNA extraction using a modified CTAB-based protocol, as previously described [21]. The first quality control of the DNA was a NanoDrop measurement, followed by agarose gel electrophoresis. After obtaining a suitable sample, a Qubit measurement was performed for accurate quantification, as previously described [20]. Afterwards, short DNA fragments were depleted with the Short Read Eliminator kit (Pacific Biosciences) following the supplier’s instructions.

Libraries for the nanopore sequencing were prepared with the SQK-LSK114 ligation-based kit (Oxford Nanopore Technologies) using 1 µg of DNA and following the supplier’s instructions. Sequencing was conducted on a PromethION 2 Solo with R10.4.1 flow cells on the MinKNOW software v25.05.14. Upon blocking a large proportion of the nanopores, a wash step was performed, followed by loading a fresh library to achieve optimal flow cell performance. Basecalling of the raw sequencing data was performed with Dorado v1.4.0 (ONT) with the high accuracy basecalling (HAC) model (dna_r10.4.1_e8.2_400bps_hac@v5.2.0), and Dorado v2.0.1 (ONT) using the super accurate basecalling (SUP) model (dna_r10.4.1_e8.2_400bps_sup@v5.2.0), both on an NVIDIA L4 GPU in the de.NBI cloud.

### Genome sequence assembly

In total, four genome assemblies were generated: for the HAC-basecalled data, assemblies were performed with Shasta [22] and Hifiasm-0.25.0-r726 [23]. For the SUP-basecalled data, Hifiasm was applied. To improve the contiguity of the representative genome sequence, the Hifiasm assembly generated from SUP-basecalled reads was merged with the Shasta assembly generated from HAC-basecalled reads. Whole-genome alignment between the two assemblies was performed with nucmer from the MUMmer4 package [24], using a minimum alignment length of 100 bp. Resulting alignments were filtered with delta-filter (MUMmer4), retaining only alignments with ≥95% identity and applying reciprocal best-hit filtering (-r -q). The filtered alignment was used to merge the two assemblies with quickmerge v0.3 [25], using the Hifiasm assembly based on SUP-basecalled reads as the self/reference assembly and the Shasta assembly as the query, with a high-confidence overlap cutoff (-hco) of 5.0 and a contig-extension cutoff (-c) of 7.0. Assembly completeness was assessed using BUSCO v6.0.0 [18] with the ‘eudicotyledons_odb12’ lineage and compared against the original three assemblies to confirm that merging improved contiguity without introducing missing or duplicated BUSCO genes. The assembly statistics were calculated using the ‘contig_stats3.py’ script [26]. Contig names were also cleaned to avoid technical issues using the ‘clean_genomic_fasta.py’ script [27].

### Genome sequence annotation

RNA-seq datasets (Additional file 1) were retrieved from the Sequence Read Archive [28] using ‘fastq-dump’ [29] to generate hints for the structural annotation. The RNA-seq read mapping was performed with HISAT2 v2.2.1 [30], and the resulting BAM file was supplied to GeMoMa v1.9 [31], enabling the inference of hints for gene prediction. The following five datasets from related species were also used as hints: *Lonicera macranthoides* (GCA_054790715.1 and GCA_054790735.1), *Lonicera japonica* [32,33]*, Lonicera japonica* (GCA_021464415.1) [34,35], and *Triplostegia glandulifera* [36,37].

The initial annotation was filtered with GeMoMa [24] using the following criteria: f=“start==’M’ and stop==’*’ and (score/aa>=’0.75’)”, the used RNA-seq evidence was prepared with GeMoMa’s ‘ERE’ based on the read mapping generated with HISAT2 [23], followed by an analysis of all predicted polypeptide sequences with BUSCO v6.0.0 [31] with the eudicotyledons_odb12 lineage, using the protein mode to assess the annotation completeness. The last filtering was done with: f=“start==’M’ and stop==’*’ and (score/aa>=’1.75’)”, and the resulting predicted genes were renamed with ‘AnnotationFinalizer’, and CDS and polypeptide sequences were extracted with GeMoMa’s ‘Extractor’.

### Functional annotation with focus on valerenic acid and flavonoid biosynthesis

Candidate genes for valerenic acid biosynthesis were identified through both sequence similarity search and domain-based genome mining. The characterized valerian terpene synthase family [9] and the cytochrome P450 VoCYP71DJ1 [10], which together constitute the entry-point and pathway-completing enzymes of valerenic acid biosynthesis, characterized through heterologous expression, were used as BLASTP queries [38] against the predicted polypeptide sequences (e-value cutoff 1e-20). The protein sequences used as baits are the following: VoTPS1, VoTPS2, VoTPS3, VoTPS4, VoTPS5, VoTPS6, VoTPS7 [9], and VoCYP71DJ1 [10]. In parallel, the predicted polypeptide sequences were screened for terpene synthase domain architecture using HMMER v3.4 [39] with the Pfam [40] terpene synthase N-terminal (PF01397) and C-terminal (PF03936) domain models; candidates matching both domains were considered high-confidence terpene synthase family members. Genome-wide identification of biosynthetic gene clusters, including terpene-type clusters, was additionally performed with plantiSMASH v2.0.4 [41].

A general prediction of gene functions was performed with the Python script ‘construct_anno.py’ [42] based on orthology to sequences in the Araport11 annotation of *Arabidopsis thaliana* [43,44]. A comprehensive annotation of the flavonoid biosynthesis genes was conducted with KIPEs v3.2.7 [45]. The anthocyanin biosynthesis-regulating MYBs were identified using MYB_annotator v1.0.3 [53], and the anthocyanin biosynthesis-regulating bHLHs were identified using bHLH_annotator v0.2 [46].

## Results and Discussion

### Chromosome counting

To confirm the chromosome number and ploidy level of the *Valeriana officinalis* samples, we analyzed at least 10 mitotic metaphases from each of the 5 available individuals. All the samples showed 28 chromosomes, corresponding to a tetraploid composition (2*n* = 4x = 28; **Figure 2**), with metacentric and submetacentric morphology.

**Figure 2.**
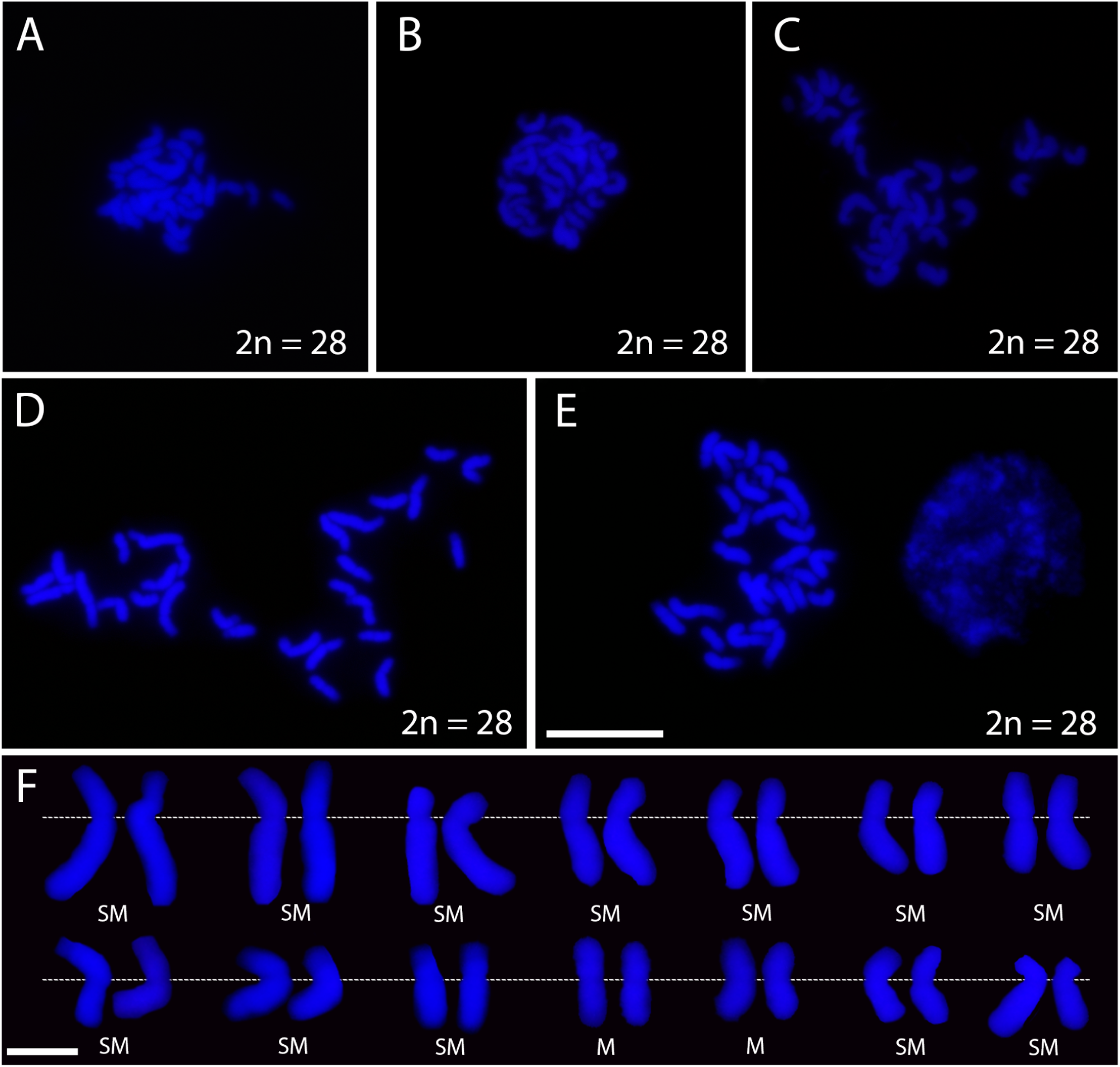
Mitotic metaphase chromosomes of *Valeriana officinalis*. **A-E:** mitotic metaphase cells of five *V. officinalis* individuals; chromosomes are counterstained with DAPI (blue). All individuals showed 2*n* = 4*x* = 28 chromosomes. **F:** Chromosome karyotype showing each chromosome pair, arranged in order from the largest to the smallest in size, based on the metaphase cell in **D**. Dashed line marks the centromeric primary constriction; SM corresponds to submetacentric chromosome pairs, M corresponds to metacentric chromosome pairs. Bar in **E** corresponds to 2.5 μm and in **F** to 1.25 μm.

### Genome sequence and annotation

Four genome sequences of *Valeriana officinalis* were assembled based on ONT long-read sequencing data. Two assemblies were generated from HAC-basecalled data, one assembly was generated from SUP-basecalled data, and one final assembly was generated by merging the SUP-basecalled Hifiasm assembly with the HAC-basecalled Shasta assembly (**Table 1**). The assembly sizes ranging from 3.1 Gbp to 3.7 Gbp are consistent with results of a flow cytometry study, which reported a DNA amount of 4.08 pg for 1C [47]. The final merged assembly comprised 153 contigs totalling 3.3 Gbp, with an N50 of 110.8 Mbp and a GC content of 37.4%, indicating a high level of contiguity suitable for comparative genomics. Assembly completeness was estimated at 97.4% based on BUSCO genes (eudicotyledons_odb12), consistent with completeness estimates for the individual input assemblies, indicating that merging improved contiguity without reducing completeness. Duplication is estimated at 97.1%, consistent with a tetraploid genome [48].

**Table 1:** Statistics of the *Valeriana officinalis* genome sequences produced in this study. A total of 2805 BUSCO genes was used for the completeness assessment.

|  | ***Shasta***<br>**(HAC)** | ***Hifiasm***<br>**(HAC)** | ***Hifiasm***<br>**(SUP)** | ***Final Assembly*** |
| --- | --- | --- | --- | --- |
| **Total assembly size** | 3.1 Gbp | 3.1 Gbp | 3.7 Gbp | 3.3 Gbp |
| **Number of sequences** | 1246 (contigs) | 1325 (contigs) | 1456 (contigs) | 153 (contigs) |
| **Maximal contig length** | 37.1 Mbp | 0.19 Mbp | 184.4 Mbp | 223.7 Mbp |
| **N50** | 6.1 Mbp | 93.3 Mbp | 3.1 Mbp | 110.8 Mbp |
| **BUSCO (complete)** | C:97.5%<br>[S:1.4%,<br>D:96.1%], | C:97.5%<br>[S:1.6%,<br>D:95.9%], | C:97.5%<br>[S:0.3%,<br>D:97.2%], | C:97.4%<br>[S:0.4%,<br>D:97.1%], |
| **BUSCO (fragmented, missing)** | F:0.7%,<br>M:1.8%,<br>E:16.5% | F:0.8%,<br>M:1.7%,<br>E:16.6% | F:0.7%,<br>M:1.7%,<br>E:15.6% | F:0.7%,<br>M:1.8%,<br>E:16.5% |

The filtered GeMoMa annotation based on hints from various related plant species (**Table 2**) emerged as the best structural annotation and was subsequently used for all downstream analyses. A completeness check of the annotation with BUSCO revealed about 96.6% of all expected genes, with 95.6% duplication, which is expected for a tetraploid genome. The total number of genes of 91,026 is comparable to the average number of genes in a tetraploid plant species [49,50] and was not considered annotation redundancy.

**Table 2:** Comparison of GeMoMa structural annotation approaches of *Valeriana officinalis*. GeMoMa was supplied with RNA-seq hints (Additional file 1) and datasets of *Lonicera macranthoides* (GCA_054790715.1 and GCA_054790735.1) and *Lonicera japonica* [33], *Lonicera japonica* (GCA_021464415.1) [34], and *Triplostegia glandulifera* [36]. The total number of BUSCO genes was 2805.

|  | GeMoMa | GeMoMa (filtered) |
| --- | --- | --- |
| **Number of genes** | 113,908 | 91,026 |
| **Number of transcripts** | 159,484 | 128,812 |
| **BUSCO**<br>**(complete)** | C:96.5%<br>[S:1.1%,D:95.4%] | C:96.6%<br>[S:1.0%,D:95.6%] |
| **BUSCO**<br>**(fragmented, missing)** | F:1.3%,<br>M:2.2% | F:1.1%,<br>M:2.2% |

### Candidate genes for valerenic acid biosynthesis

Valerenic acid biosynthesis begins with the cyclization of farnesyl diphosphate (FPP) by valerena-4,7(11)-diene synthase (also reported as valerena-1,10-diene synthase), the first committed and rate-limiting step of the pathway [9]. The resulting sesquiterpene is oxidized to valerenic acid through the combined activity of the cytochrome P450 VoCYP71DJ1, an alcohol dehydrogenase, and an aldehyde dehydrogenase (**Figure 3**), as demonstrated by heterologous reconstruction in yeast [10]. Thus, the most directly supported candidate genes in *V. officinalis* are those related to the previously characterized VoTPS enzymes and VoCYP71DJ1.

**Figure 3.**
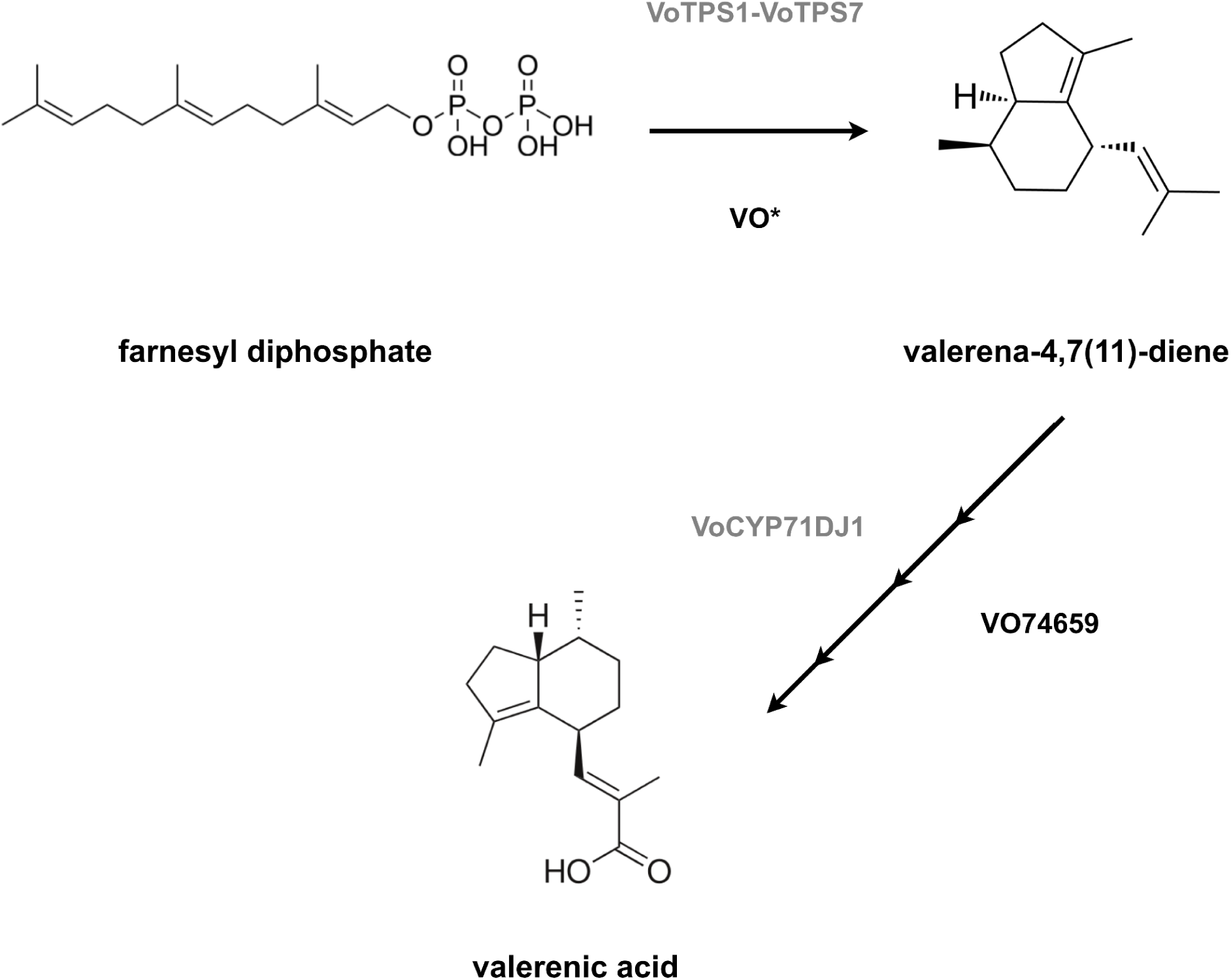
Representation of the valerenic acid biosynthesis pathway with the candidate genes identified in this study. All listed candidates carry both the TPS N-terminal (PF01397) and C-terminal (PF03936) Pfam domains. To avoid visual clutter in the image, the candidate genes for TPS were represented only by VO* and are as follows: *VO42896* (VoTPS1), *VO73413* (VoTPS2), *VO84642* (VoTPS3 and VoTPS4), *VO70249* (VoTPS5), *VO67651* (VoTPS6), and *VO31087* (VoTPS7); they can be found in Additional File 2, and the full domain-based TPS gene family can be found in Additional File 3.

Terpene synthases (TPSs) are generally characterized by a conserved two-domain architecture, an N-terminal domain (Pfam PF01397) and a metal-binding C-terminal domain (Pfam PF03936), and typically form multigene families of several dozen to over a hundred members per angiosperm genome, classified into clades TPS-a through TPS-h [11,12]. The *V. officinalis* genome sequence revealed a terpene synthase gene family of 244 loci encoding polypeptides with the complete, canonical two-domain architecture, a family size within the range reported for other sequenced angiosperm genomes [12]. All seven previously characterized VoTPS genes are represented in this family, encoding near-identical polypeptide sequences to their originally cloned sequences (99.4–100%; Additional File 2), consistent with these reference sequences having been cloned from *V. officinalis* itself [9]. Notably, VoTPS3 and VoTPS4 were proposed to be the same gene in the original characterization study [9]; consistent with this, both reference sequences recovered the identical best-matching locus (*VO84642*) in the *V. officinalis* genome, independently supporting their proposed redundancy at the genomic level. The locus encoding the pathway-completing cytochrome P450, VoCYP71DJ1, is more divergent from its reference sequence (76.3–76.5% identity; Additional File 2) than the TPS loci, mirroring the generally higher sequence divergence observed among plant cytochrome P450 paralogs even within the same CYP71 clan [51,52].

Genome-wide analysis further identified 162 candidate biosynthetic gene clusters, including 25 classified as terpene-associated and 5 clusters that combine a terpene synthase with a cytochrome P450 (Additional File 4). Among these, cluster 99 (ptg000183l:6,093,600–6,123,086; 29.5 kb, **Figure 4**) stands out as the most compact, containing only the N-terminal and C-terminal terpene synthase domains (PF01397/PF03936) and a P450 domain, this two-enzyme composition mirrors the core architecture of several experimentally characterized plant terpenoid biosynthetic gene clusters, in which a single terpene synthase is paired with one or a small number of dedicated cytochrome P450 oxidases to convert a cyclized terpene scaffold into an oxidized end product [53,54]. It is nonetheless smaller than most experimentally validated plant metabolic gene clusters, which typically range from approximately 35 kb up to several hundred kb [55].

**Figure 4.**
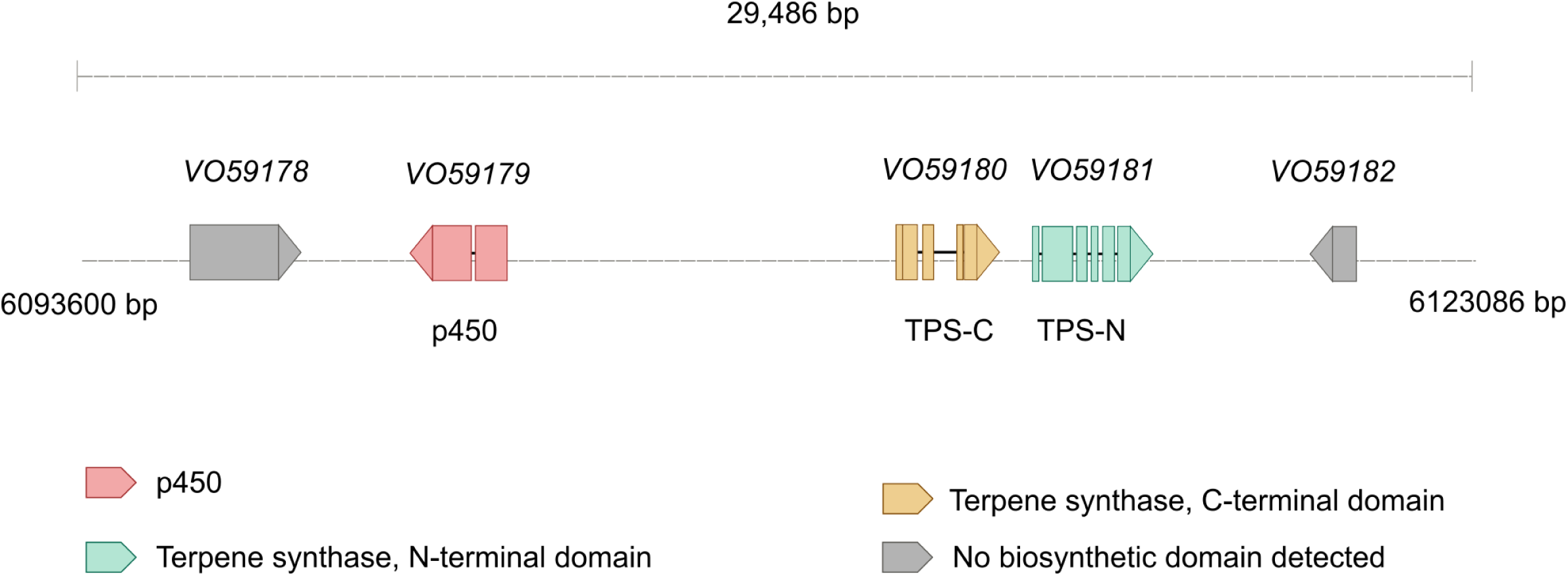
Genomic organization of cluster 99, a candidate biosynthetic gene cluster for terpenoid tailoring reactions (scaffold ptg000183: 6093600 - 6123086). The cluster is located approximately 6.1 Mbp upstream and 95.0 Mbp downstream of its host scaffold (101.1 Mbp total length). Exons are colored by detected domain: red, cytochrome P450; yellow, terpene synthase C-terminal domain; green, terpene synthase N-terminal domain; gray, no biosynthetic domain detected; thin black lines connecting exons represent introns. *VO59179*, *VO59180*, and *VO59181* represent candidate tailoring and cyclization enzymes co-localized within a single 29.5 kb cluster.

Cluster classification does not prove that a locus is involved in valerenic acid production. In particular, the presence of a TPS gene within a predicted cluster does not establish its substrate specificity, product profile, or participation in the valerenic acid pathway. Still, the recurrent co-localization of genes encoding these two enzyme classes in the *V. officinalis* genome raises the possibility that the valerenic acid pathway, or a related sesquiterpenoid pathway, is organized as such a cluster, since physical clustering of genes for specialized metabolite biosynthesis, including terpenoid pathways, is a documented feature of plant genomes, thought to facilitate co-inheritance and coordinated transcriptional regulation of complete pathways [54].

### Flavonoid biosynthesis in *Valeriana officinalis*

The core flavonoid pathway is well conserved across flowering plants, with phenylalanine-derived intermediates channeled via CHS, CHI, and F3H into dihydroflavonols, which are then partitioned toward flavonols and anthocyanins/proanthocyanidins by FLS and DFR, respectively [56]. Using KIPEs, candidates for each of these core steps in the *V. officinalis* annotation were identified (**Figure 5**). Candidate genes for all core steps in flavonol and anthocyanin biosynthesis were discovered, which is congruent with previous reports about the presence of flavonols and anthocyanins in *Valeriana* species [13]. The biosynthesis of anthocyanins can be observed visually as pink flower color (**Figure 1B**).

**Figure 5.**
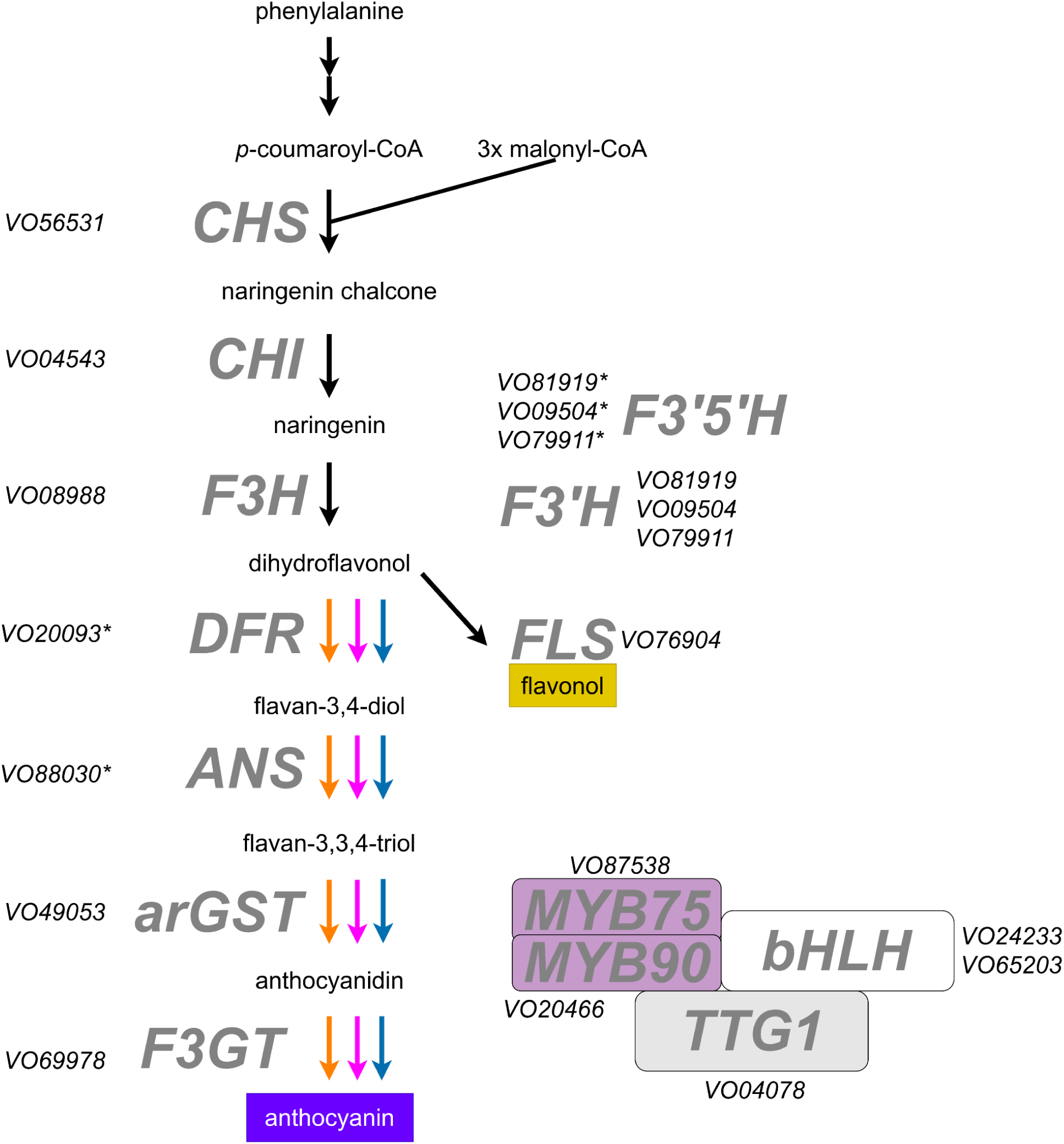
Representation of at least one structural gene of each step in the anthocyanin biosynthesis. Candidates lacking an important amino acid residue are marked with an asterisk. CHS, chalcone synthase; CHI, chalcone isomerase; F3H, flavanone 3-hydroxylase; F3’H, flavonoid 3’-hydroxylase; F3’5’H, flavonoid 3’,5’-hydroxylase; FLS, flavonol synthase; DFR, dihydroflavonol 4-reductase; ANS, anthocyanidin synthase; arGST, anthocyanin-related glutathione S-transferase; and F3GT, flavonoid 3-O-glucosyltransferase. MYB75 and MYB90, R2R3-MYB transcription factors controlling flavonoid biosynthesis; bHLH, basic helix–loop–helix transcription factor; and TTG1, TRANSPARENT TESTA GLABRA 1, a WD40-repeat protein involved in the MBW regulatory complex. More detailed gene descriptions can be found in Additional File 5.

The CHS (*VO56531*), CHI (*VO04543*), F3H (*VO08988*), FLS (*VO76904*), F3GT (*VO69978*), and arGST (*VO49053*) sequences show 100% of the amino acid residues required for functional enzymes. In contrast, the best candidates for DFR (*VO20093*) and ANS (*VO88030*) retain 96.4% and 95.7% of the expected residues, respectively. Therefore, they can be considered strong candidates, and lineage-specific differences in DFR should be explored through enzyme characterization or complementation studies in the future.

Three paralogous F3′H candidates (*VO81919*, *VO09504*, *VO79911*) were identified, each showing 100% conservation against the F3′H residue set, the same three genes also scored 98% against the F3′5′H bait set (Additional File 5), illustrating the documented difficulty of separating these two CYP75-family paralogs by sequence homology alone, since their substrate specificity is governed by a small number of active site residues [57]. Across all steps, several paralogs were recovered per gene family (Additional File 5), reflecting both the expected size of these gene families in flowering plants and the additional gene dosage harboured by the tetraploid genome of *V. officinalis*.

In total, 403 bHLH candidates and 324 MYB-domain proteins (predominantly R2R3) were identified in the *V. officinalis* annotation. For the flavonol, anthocyanin, and proanthocyanidin regulatory subgroups, several *V. officinalis* genes were recovered in proximity to the SG5 (proanthocyanidin activator), SG6 (anthocyanin activator), and SG7 (flavonol activator) reference subgroups (Additional File 6). The TT8 lineage was represented, with two loci (*VO24233*, *VO65203*) as candidates. SG6 MYBs and TT8 form a transcription factor complex with a WD40, which is known as the MBW complex and activates the anthocyanin biosynthesis genes [58]. In summary, these results indicate that *Valeriana officinalis* possesses the MYB and bHLH regulators required for the activation of the anthocyanin biosynthesis and the structural genes for the formation of pigments.

## Conclusions

This study presents the first genome sequence and annotation of *Valeriana officinalis*, filling an important gap for this medicinal plant. We present a genome sequence of high contiguity and completeness. The downstream analysis, in particular, the identification of candidate loci for valerenic acid biosynthesis and genes related to the flavonoid pathway, offers valuable starting points for investigating how these compounds are produced and regulated in the native plant. Overall, this resource should accelerate efforts to understand valerian metabolism and may ultimately support breeding or biotechnological projects towards enhanced production of therapeutically important compounds.

## Data availability

All data sets underlying this study are publicly available. Sequencing data have been deposited at the European Nucleotide Archive (PRJEB115730). The assembled genome sequence and corresponding annotation are available via bonndata (https://doi.org/10.60507/FK2/TQZVWU).

## Acknowledgements

This work was supported by the de.NBI Cloud within the German Network for Bioinformatics Infrastructure (de.NBI) and ELIXIR-DE (Forschungszentrum Jülich and W-de.NBI-001, W-de.NBI-004, W-de.NBI-008, W-de.NBI-010, W-de.NBI-013, W-de.NBI-014, W-de.NBI-016, W-de.NBI-022). We thank all members of the research group Plant Biotechnology and Bioinformatics for their discussion and support. We acknowledge support from Project DEAL and the University of Bonn for open access publication. We are grateful for the excellent support provided by the team of the University of Bonn Botanic Gardens.

## Author contributions

JAVSdO and BP designed the experiment. JAVSdO did the sequencing. MAB performed the chromosome identification and karyotype analysis. JAVSdO conducted the bioinformatic analyses. JAVSdO wrote the manuscript with input from all authors. All authors reviewed the final version of the manuscript and consented to its submission.

## Additional Files

Additional file 1: RNA-seq datasets used as hints for structural annotation.

Additional file 2: List of TPS candidate genes.

Additional file 3: List of full domain-based TPS candidates.

Additional file 4: Biosynthetic gene cluster predictions.

Additional file 5: All candidate genes associated with the flavonoid biosynthesis pathway.

Additional file 6: Gene tree of *Valeriana* MYB candidates associated with flavonoid biosynthesis in context with previously characterized MYBs.

## Notes

https://doi.org/10.60507/FK2/TQZVWU

### Competing Interest Statement

The authors have declared no competing interest.

## References

1. Bressler S, Klatte-Asselmeyer V, Fischer A, Paule J, Dobeš C. Variation in genome size in the Valeriana officinalis complex resulting from multiple chromosomal evolutionary processes. Preslia. 2017;89:41–61. 10.23855/preslia.2017.041

2. Plant DNA C-values Database | Royal Botanic Gardens, Kew. https://cvalues.science.kew.org/. Accessed 26 Mar 2026

3. Sezen Karaoğlan E. Valeriana officinalis L. In: Gürağaç Dereli FT, Ilhan M, Belwal T, editors. Nov Drug Targets Tradit Herb Med Sci Clin Evid. Cham: Springer International Publishing; 2022. p. 565–8. 10.1007/978-3-031-07753-1_37

4. Shinjyo N, Waddell G, Green J. Valerian Root in Treating Sleep Problems and Associated Disorders—A Systematic Review and Meta-Analysis. J Evid-Based Integr Med. SAGE Publications Inc STM; 2020;25:2515690X20967323. 10.1177/2515690X20967323

5. Anxiety disorders. https://www.who.int/news-room/fact-sheets/detail/anxiety-disorders. Accessed 24 July 2026

6. Hattesohl M, Feistel B, Sievers H, Lehnfeld R, Hegger M, Winterhoff H. Extracts of *Valeriana officinalis* L. s.l. show anxiolytic and antidepressant effects but neither sedative nor myorelaxant properties. Phytomedicine. 2008;15:2–15. 10.1016/j.phymed.2007.11.027

7. Murphy K, Kubin ZJ, Shepherd JN, Ettinger RH. *Valeriana officinalis* root extracts have potent anxiolytic effects in laboratory rats. Phytomedicine. 2010;17:674–8. 10.1016/j.phymed.2009.10.020

8. Becker A, Felgentreff F, Schröder H, Meier B, Brattström A. The anxiolytic effects of a Valerian extract is based on Valerenic acid. BMC Complement Altern Med. 2014;14:267. 10.1186/1472-6882-14-267

9. Yeo Y-S, Nybo SE, Chittiboyina AG, Weerasooriya AD, Wang Y-H, Góngora-Castillo E, et al. Functional Identification of Valerena-1,10-diene Synthase, a Terpene Synthase Catalyzing a Unique Chemical Cascade in the Biosynthesis of Biologically Active Sesquiterpenes in *Valeriana officinalis*\*. J Biol Chem. 2013;288:3163–73. 10.1074/jbc.M112.415836

10. Wong J, d’Espaux L, Dev I, van der Horst C, Keasling J. *De novo* synthesis of the sedative valerenic acid in *Saccharomyces cerevisiae*. Metab Eng. 2018;47:94–101. 10.1016/j.ymben.2018.03.005

11. Bohlmann J, Meyer-Gauen G, Croteau R. Plant terpenoid synthases: Molecular biology and phylogenetic analysis. Proc Natl Acad Sci. Proceedings of the National Academy of Sciences; 1998;95:4126–33. 10.1073/pnas.95.8.4126

12. Chen F, Tholl D, Bohlmann J, Pichersky E. The family of terpene synthases in plants: a mid-size family of genes for specialized metabolism that is highly diversified throughout the kingdom. Plant J. 2011;66:212–29. 10.1111/j.1365-313X.2011.04520.x

13. Navarrete A, Avula B, Choi Y-W, Khan IA. Chemical Fingerprinting of Valeriana Species: Simultaneous Determination of Valerenic Acids, Flavonoids, and Phenylpropanoids Using Liquid Chromatography with Ultraviolet Detection. J AOAC Int. 2006;89:8–15. 10.1093/jaoac/89.1.8

14. Średnicka-Tober D, Hallmann E, Kopczyńska K, Góralska-Walczak R, Barański M, Grycz A, et al. Profile of Selected Secondary Metabolites and Antioxidant Activity of Valerian and Lovage Grown in Organic and Low-Input Conventional System. Metabolites. Multidisciplinary Digital Publishing Institute; 2022;12:835. 10.3390/metabo12090835

15. Winkel-Shirley B. Flavonoid Biosynthesis. A Colorful Model for Genetics, Biochemistry, Cell Biology, and Biotechnology. Plant Physiol. 2001;126:485–93. 10.1104/pp.126.2.485

16. Grünig N, Horz JM, Pucker B. Diversity and ecological functions of anthocyanins. BMC Plant Biol. 2025;26:146. 10.1186/s12870-025-08006-3

17. Butelli E, Titta L, Giorgio M, Mock H-P, Matros A, Peterek S, et al. Enrichment of tomato fruit with health-promoting anthocyanins by expression of select transcription factors. Nat Biotechnol. Nature Publishing Group; 2008;26:1301–8. 10.1038/nbt.1506

18. Appelhagen I, Wulff-Vester AK, Wendell M, Hvoslef-Eide A-K, Russell J, Oertel A, et al. Colour bio-factories: Towards scale-up production of anthocyanins in plant cell cultures. Metab Eng. 2018;48:218–32. 10.1016/j.ymben.2018.06.004

19. Choudhary N, Khatun N, Pucker B. Almost 200 Years of Anthocyanin Research: What We Know, What We Assume, and What Remains Unknown. Preprints; 2026. 10.20944/preprints202603.2058.v1

20. de Oliveira JAVS, Choudhary N, Meckoni SN, Nowak MS, Hagedorn M, Pucker B. Cookbook for plant genome sequences. BMC Genomics. 2026;27:231. 10.1186/s12864-026-12623-z

21. Siadjeu C, Pucker B, Viehöver P, Albach DC, Weisshaar B. High Contiguity de novo Genome Sequence Assembly of Trifoliate Yam (Dioscorea dumetorum) Using Long Read Sequencing. Genes. Multidisciplinary Digital Publishing Institute; 2020;11:274. 10.3390/genes11030274

22. Shafin K, Pesout T, Lorig-Roach R, Haukness M, Olsen HE, Bosworth C, et al. Nanopore sequencing and the Shasta toolkit enable efficient de novo assembly of eleven human genomes. Nat Biotechnol. Nature Publishing Group; 2020;38:1044–53. 10.1038/s41587-020-0503-6

23. Cheng H, Concepcion GT, Feng X, Zhang H, Li H. Haplotype-resolved de novo assembly using phased assembly graphs with hifiasm. Nat Methods. Nature Publishing Group; 2021;18:170–5. 10.1038/s41592-020-01056-5

24. Marçais G, Delcher AL, Phillippy AM, Coston R, Salzberg SL, Zimin A. MUMmer4: A fast and versatile genome alignment system. PLOS Comput Biol. Public Library of Science; 2018;14:e1005944. 10.1371/journal.pcbi.1005944

25. Chakraborty M, Baldwin-Brown JG, Long AD, Emerson JJ. Contiguous and accurate de novo assembly of metazoan genomes with modest long read coverage. Nucleic Acids Res. 2016;44:e147. 10.1093/nar/gkw654

26. Meckoni SN, Nass B, Pucker B. Phylogenetic placement of Ceratophyllum submersum based on a complete plastome sequence derived from nanopore long read sequencing data. BMC Res Notes. 2023;16:187. 10.1186/s13104-023-06459-z

27. Pucker B. bpucker/GenomeAssembly. 2026. https://github.com/bpucker/GenomeAssembly. Accessed 24 July 2026

28. Leinonen R, Sugawara H, Shumway M, on behalf of the International Nucleotide Sequence Database Collaboration. The Sequence Read Archive. Nucleic Acids Res. 2011;39:D19–21. 10.1093/nar/gkq1019

29. ncbi/sra-tools. NCBI - National Center for Biotechnology Information/NLM/NIH; 2026. https://github.com/ncbi/sra-tools. Accessed 24 July 2026

30. Kim D, Paggi JM, Park C, Bennett C, Salzberg SL. Graph-based genome alignment and genotyping with HISAT2 and HISAT-genotype. Nat Biotechnol. Nature Publishing Group; 2019;37:907–15. 10.1038/s41587-019-0201-4

31. Keilwagen J, Wenk M, Erickson JL, Schattat MH, Grau J, Hartung F. Using intron position conservation for homology-based gene prediction. Nucleic Acids Res. 2016;44:e89. 10.1093/nar/gkw092

32. Genome Warehouse. https://ngdc.cncb.ac.cn/gwh/Assembly/660/show. Accessed 24 July 2026

33. Pu X, Li Z, Tian Y, Gao R, Hao L, Hu Y, et al. The honeysuckle genome provides insight into the molecular mechanism of carotenoid metabolism underlying dynamic flower coloration. New Phytol. 2020;227:930–43. 10.1111/nph.16552

34. Yu H, Guo K, Lai K, Shah MA, Xu Z, Cui N, et al. Chromosome-scale genome assembly of an important medicinal plant honeysuckle. Sci Data. Nature Publishing Group; 2022;9:226. 10.1038/s41597-022-01385-4

35. Genome annotation for Sijihua, which is a stress-resistance honeysuckle variety. figshare. figshare; 2022. 10.6084/m9.figshare.18092708.v6

36. Zhang J, Dong K-L, Ren M-Z, Wang Z-W, Li J-H, Sun W-J, et al. Coping with alpine habitats: genomic insights into the adaptation strategies of Triplostegia glandulifera (Caprifoliaceae). Hortic Res. 2024;11:uhae077. 10.1093/hr/uhae077

37. Zhang J, Sun W. Triplostegia glandulifera genome assembly and annotation. 2024. 10.6084/m9.figshare.25018103

38. Camacho C, Coulouris G, Avagyan V, Ma N, Papadopoulos J, Bealer K, et al. BLAST+: architecture and applications. BMC Bioinformatics. 2009;10:421. 10.1186/1471-2105-10-421

39. Eddy SR. Accelerated Profile HMM Searches. PLOS Comput Biol. Public Library of Science; 2011;7:e1002195. 10.1371/journal.pcbi.1002195

40. Mistry J, Chuguransky S, Williams L, Qureshi M, Salazar GA, Sonnhammer ELL, et al. Pfam: The protein families database in 2021. Nucleic Acids Res. 2021;49:D412–9. 10.1093/nar/gkaa913

41. Kautsar SA, Suarez Duran HG, Blin K, Osbourn A, Medema MH. plantiSMASH: automated identification, annotation and expression analysis of plant biosynthetic gene clusters. Nucleic Acids Res. 2017;45:W55–63. 10.1093/nar/gkx305

42. Pucker B, Iorizzo M. Apiaceae FNS I originated from F3H through tandem gene duplication. PLOS ONE. Public Library of Science; 2023;18:e0280155. 10.1371/journal.pone.0280155

43. Cheng C-Y, Krishnakumar V, Chan AP, Thibaud-Nissen F, Schobel S, Town CD. Araport11: a complete reannotation of the Arabidopsis thaliana reference genome. Plant J. 2017;89:789–804. 10.1111/tpj.13415

44. Lamesch P, Berardini TZ, Li D, Swarbreck D, Wilks C, Sasidharan R, et al. The Arabidopsis Information Resource (TAIR): improved gene annotation and new tools. Nucleic Acids Res. 2012;40:D1202–10. 10.1093/nar/gkr1090

45. Rempel A, Choudhary N, Pucker B. KIPEs3: Automatic annotation of biosynthesis pathways. PLOS ONE. Public Library of Science; 2023;18:e0294342. 10.1371/journal.pone.0294342

46. Thoben C, Pucker B. Automatic annotation of the bHLH gene family in plants. BMC Genomics. 2023;24:780. 10.1186/s12864-023-09877-2

47. Hidalgo O, Mathez J, Garcia S, Garnatje T, Pellicer J, Vallès J. Genome Size Study in the Valerianaceae: First Results and New Hypotheses. J Bot. 2010;2010:797246. 10.1155/2010/797246

48. Wolff K, Nowak MS, Thoben C, Beuerle T, Pucker B. Rubus armeniacus genome sequence reveals the secrets of blackberry anthocyanin biosynthesis. bioRxiv; 2026. p. 2026.05.05.723051. 10.64898/2026.05.05.723051

49. Wang F, Xia Z, Zou M, Zhao L, Jiang S, Zhou Y, et al. The autotetraploid potato genome provides insights into highly heterozygous species. Plant Biotechnol J. 2022;20:1996–2005. 10.1111/pbi.13883

50. Yim WC, Swain ML, Ma D, An H, Bird KA, Curdie DD, et al. The final piece of the Triangle of U: Evolution of the tetraploid Brassica carinata genome. Plant Cell. 2022;34:4143–72. 10.1093/plcell/koac249

51. Nelson D, Werck-Reichhart D. A P450-centric view of plant evolution. Plant J. 2011;66:194–211. 10.1111/j.1365-313X.2011.04529.x

52. Mizutani M, Ohta D. Diversification of P450 Genes During Land Plant Evolution. Annu Rev Plant Biol. Annual Reviews; 2010;61:291–315. 10.1146/annurev-arplant-042809-112305

53. Field B, Osbourn AE. Metabolic Diversification—Independent Assembly of Operon-Like Gene Clusters in Different Plants. Science. American Association for the Advancement of Science; 2008;320:543–7. 10.1126/science.1154990

54. Nützmann H-W, Osbourn A. Gene clustering in plant specialized metabolism. Curr Opin Biotechnol. 2014;26:91–9. 10.1016/j.copbio.2013.10.009

55. Bharadwaj R, Kumar SR, Sharma A, Sathishkumar R. Plant Metabolic Gene Clusters: Evolution, Organization, and Their Applications in Synthetic Biology. Front Plant Sci. Frontiers; 2021;12. 10.3389/fpls.2021.697318

56. Pucker B, Selmar D. Biochemistry and Molecular Basis of Intracellular Flavonoid Transport in Plants. Plants. 2022;11:963. 10.3390/plants11070963

57. Seitz C, Ameres S, Forkmann G. Identification of the molecular basis for the functional difference between flavonoid 3′-hydroxylase and flavonoid 3′,5′-hydroxylase. FEBS Lett. 2007;581:3429–34. 10.1016/j.febslet.2007.06.045

58. Gonzalez A, Zhao M, Leavitt JM, Lloyd AM. Regulation of the anthocyanin biosynthetic pathway by the TTG1/bHLH/Myb transcriptional complex in Arabidopsis seedlings. Plant J. 2008;53:814–27. 10.1111/j.1365-313X.2007.03373.x

