## Supplementary figures and images for "*Valeriana officinalis* genome sequence reveals candidate genes for valerenic acid biosynthesis and flavonoid metabolism"

### Additional File 6

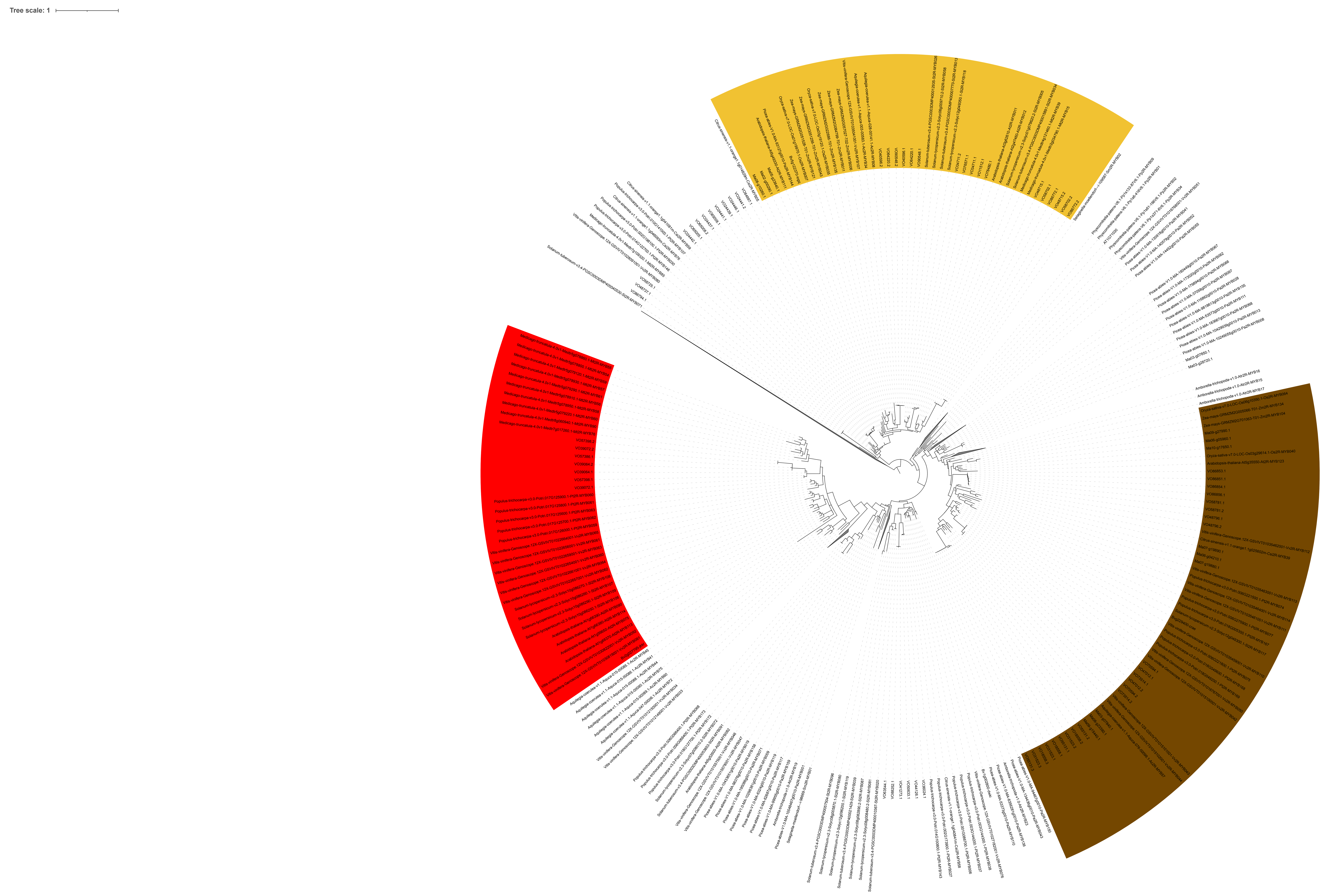
